# Gender (im)balance in citation practices in cognitive neuroscience

**DOI:** 10.1101/2020.08.19.257402

**Authors:** Jacqueline M. Fulvio, Ileri Akinnola, Bradley R. Postle

## Abstract

In the field of neuroscience, despite the fact that the proportion of peer-reviewed publications authored by women has increased in recent decades, the proportion of citations of women-led publications has not seen a commensurate increase: In five broad-scope journals, citations of papers first- and/or last-authored by women have been shown to be fewer than would be expected if gender was not a factor in citation decisions (Dworkin et al., 2020). Given the important implications that such underrepresentation may have on the careers of women researchers, it is important to determine whether this same trend is true in subdisciplines of the field, where interventions might be more effective. Here, we report the results of an extension of the analyses carried out by Dworkin et al. (2020) to citation patterns in the Journal of Cognitive Neuroscience (*JoCN*). The results indicate that the underrepresentation of women-led publications in reference sections is also characteristic of papers published in *JoCN* over the past decade. Furthermore, this pattern of citation imbalances is present for all gender classes of authors, implicating systemic factors. These results contribute to the growing body of evidence that intentional action is needed to address inequities in the way that we carry out and communicate our science.

## INTRODUCTION

The public dissemination of research findings is critical for the advancement of any field of scientific inquiry. Similarly, evidence of impactful publication in peer-reviewed journals is critical for a researcher’s advancement in their field. For example, citation-based metrics such as impact factors, the h-Index (Hirsch, 2005), and the i10-index (Connor, 2011) contribute to the evaluation of one’s scholarly “worth” (Fairhall & Marder, 2020). It is problematic, therefore, that citation-based metrics of scholarship in neuroscience show a gender bias. A recent study evaluating citation practices in five broad-scope neuroscience journals – *Brain*, the *Journal of Neuroscience*, *Nature Neuroscience*, *NeuroImage*, and *Neuron* – demonstrated over-citation of articles published by men as first and last authors compared to the rate expected if gender did not play a role in citation choices, whereas articles published by at least one woman in the first-or last-author position have been under-cited (Dworkin et al., 2020).

The findings of Dworkin et al. (2020) are a cautionary tale for fields grappling with gender disparities because bias in citation practices may limit the advancement of individual researchers, as well as advancement of their approaches and ideas. Quantification and dissemination of evidence of such biases is an important first step toward developing more equitable practices. Here, we sought to determine whether the gender imbalance in citation practices reported for broad-scope neuroscience journals is also characteristic of the *Journal of Cognitive Neuroscience* (*JoCN*), the flagship journal of this subdiscipline of neuroscience.

## METHODS

We applied the methodological approach used by Dworkin et al. (2020), using their open-source R code (https://osf.io/h79g8/). Where necessary, we modified the code to support the *JoCN*-specific analysis.

### Data acquisition

The data for the analysis were obtained from the Web of Science website (https://www.webofknowledge.com/). Metadata for the 2,106 research articles and review papers published in *JoCN* from January 2009-July 2020 were downloaded. We note that metadata for *JoCN* articles are available dating back to 1995, but metadata from prior to 2009 contain author initials rather than full first names, the latter being necessary for the analysis. Additionally, pre-processed metadata from broad-scope neuroscience journals – *Brain*, the *Journal of Neuroscience*, *Nature Neuroscience*, *NeuroImage*, and *Neuron* (from here forward, the “broad-scope journals”) -- were obtained from Jordan Dworkin with permission for use in the analysis described here.

### Gender category assignment

#### Gender category assignment of papers published in JoCN

We first extracted the names of each author of each publication from the metadata into an array with the general format, “last name, first name;last name, first name”. We then implemented the algorithm used by Dworkin et al. (2020) to disambiguate authors with different versions of their name across papers (such as instances with and without middle initials, or with and without nicknames). In brief, the algorithm matches entries first by last name and then by the same first and/or middle name or initials and assigns the most common first name variant to all instances.

Next, the first names of the first and last authors of each paper were assigned a probability of belonging to someone self-identifying with either of two gender labels -- ‘man’ or ‘woman’. (Note that the a priori assumption that gender identification is a binary variable is not correct but was necessitated by limitations of our method.) First, each name was queried within the Social Security Administration (SSA) baby name dataset, which assigns labels based on the sex assigned at birth. If the name was found, the probabilities of that name belonging to someone self-identifying as ‘man’ and ‘woman’ were returned. If the name was not found, it was submitted to Gender API (http://gender-api.com/) for probability assignment. Gender API includes approximately 815,000 unique first names from 189 countries and assigns labels based on a combination of the sex assigned at birth and genders detected in social media profiles. Using the same criteria as Dion et al. (2018) and Dworkin et al. (2020), we assigned a gender label to each author if their name had a probability ≥0.70 of belonging to someone of either gender.

The 2,106 research articles published in *JoCN* were then assigned an authorship gender category (‘man/man’ (MM), ‘woman/man’ (WM), ‘man/woman’ (MW), or ‘woman/woman’ (WW)) based on the assigned gender labels of the first and last authors. Upon completion of this step, ~9% of the articles had incomplete authorship gender category designations due to single authorship, or middle or first name initials, or other formatting problems that arose during metadata extraction that impeded the name query steps. In these cases, we performed a manual correction step to hand-code the category designations (Dion et al., 2018), sometimes entailing visits to the article’s page on the journal’s website, or the author’s website. Single-authored papers were assigned ‘MM’ or ‘WW’ according to the assigned gender label of the author.

#### Gender category assignment of papers cited in papers published in JoCN

A final critical pre-processing step was to assign authorship gender categories to the papers cited in the 2,106 *JoCN* articles. We first extracted the citation lists from the metadata of the 2,106 articles. Importantly, although the citation lists did not contain author first names (only initials), they did contain the Digital Object Identifier (DOI) of each citation. These DOIs allowed us to use the pre-processed data from six journals – the five broad-scope journals plus *JoCN* -- as a lookup table, in which we attempted to match each citation DOI with a DOI from one of the six journals. If a match was found, we assigned that cited paper to the authorship gender category assigned to the original article (from the previous step for *JoCN*, and from analogous data for the five broad-scope journals analyzed by Dworkin et al. (2020).

### Categorical gender quantification

#### Quantification of JoCN authorship

The gender balance of authorship in *JoCN* was quantified in three ways: collapsing across the January 2009-July 2020 time frame; broken out into each publication month; and as the cumulative portion of the overall time frame leading up to each publication month and year. This latter set of proportions served as the base rate (i.e., the *expected* proportions) for the Gender Citation Balance Index calculations described below.

#### Computation of Gender Citation Balance Indices

Our primary goal was to determine how the gender proportions in the reference lists of papers published in *JoCN* corresponded to the gender proportions of *JoCN* authorship. Following Dworkin et al. (2020), we removed self-citations (i.e., cited articles for which either the first or last author was the either first or last author on the citing paper) to remove effects of gender differences in self-citation behaviors (King et al., 2017), and focused instead on authors’ citation of other researchers in the field. For each of the 2,069 articles that cited other articles from *JoCN* and the five broad-scope journals, we computed the proportion of those citations assigned to each of the four gender citation categories, which were designated the *observed* proportions.

We computed Gender Citation Balance Indices for each of the four gender citation categories as:

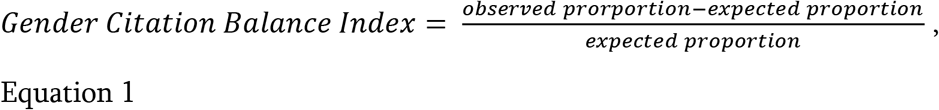

Thus, positive values corresponded to more frequent citations of the category than expected, and negative values corresponded to less frequent citations of the category than expected. (Note that because we did not have *JoCN* publication data prior to January 2009, we used the *JoCN* authorship in each category during January 2009 for the expected rates of articles published in January 2009.) Finally, we bootstrapped the 95% confidence interval for each category using 1000 iterations of random sampling with replacement from the 2,069 articles. For each iteration, we determined the Gender Citation Balance Index for each category, and the 2.5th and 97.5th percentiles corresponded to the lower and upper bounds of the confidence interval for that category, respectively.

### RESULTS

The categorical gender breakdown in *JoCN* authorship has been relatively stable from 2009 to mid-2020, with an increase in WW-authored publications in the most recent years (**Figure 1**). Overall, 40.8% of *JoCN* articles published during this timeframe were MM, with the remaining 59.2% having at least one woman in the first and last author positions (i.e., W∪W).

**Figure 1.**
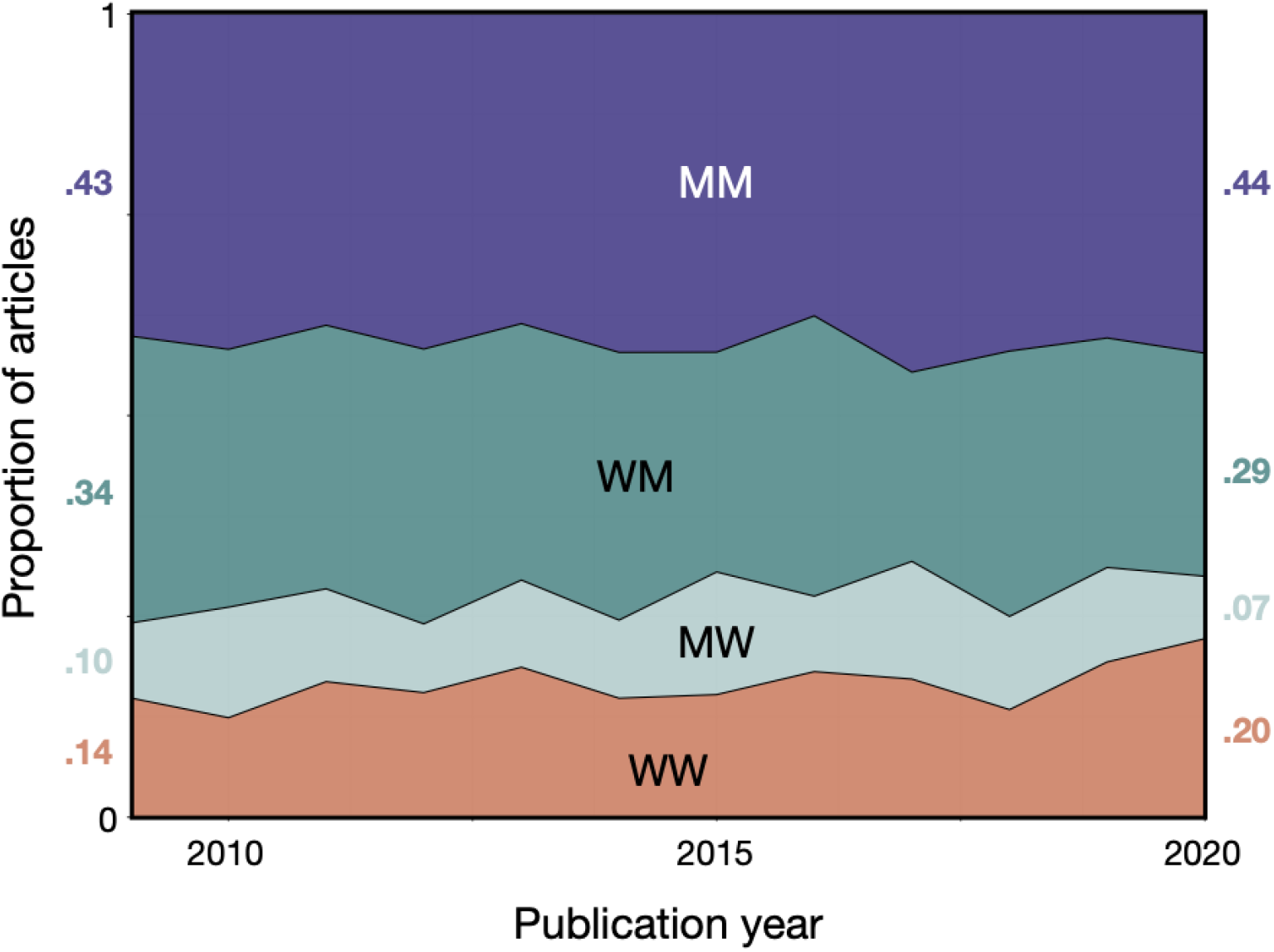
Gender breakdown in *JoCN* authorship from 2009-2020. Proportion of *JoCN* articles assigned to four categories: men as first and last author (MM; purple), women as first author and men as last author (WM; darker green), men as first author and women as last author (MW; lighter green) and women as both first and last author (WW; salmon). For ease of comparison across time, the proportions of each category are indicated for 2009 (left) and 2020 (right).

The Gender Citation Balance Indices of articles published in *JoCN* reveal an over-citation of MM papers and an under-citation of WM, MW, and WW papers (**Figure 2**). Importantly, this qualitative pattern is observed across author “gender subgroupings” when papers are broken out by author gender category, although MW papers have a positive Gender Citation Balance Index for papers from the MW and WW gender subgroups (**Figure 3**).

**Figure 2.**
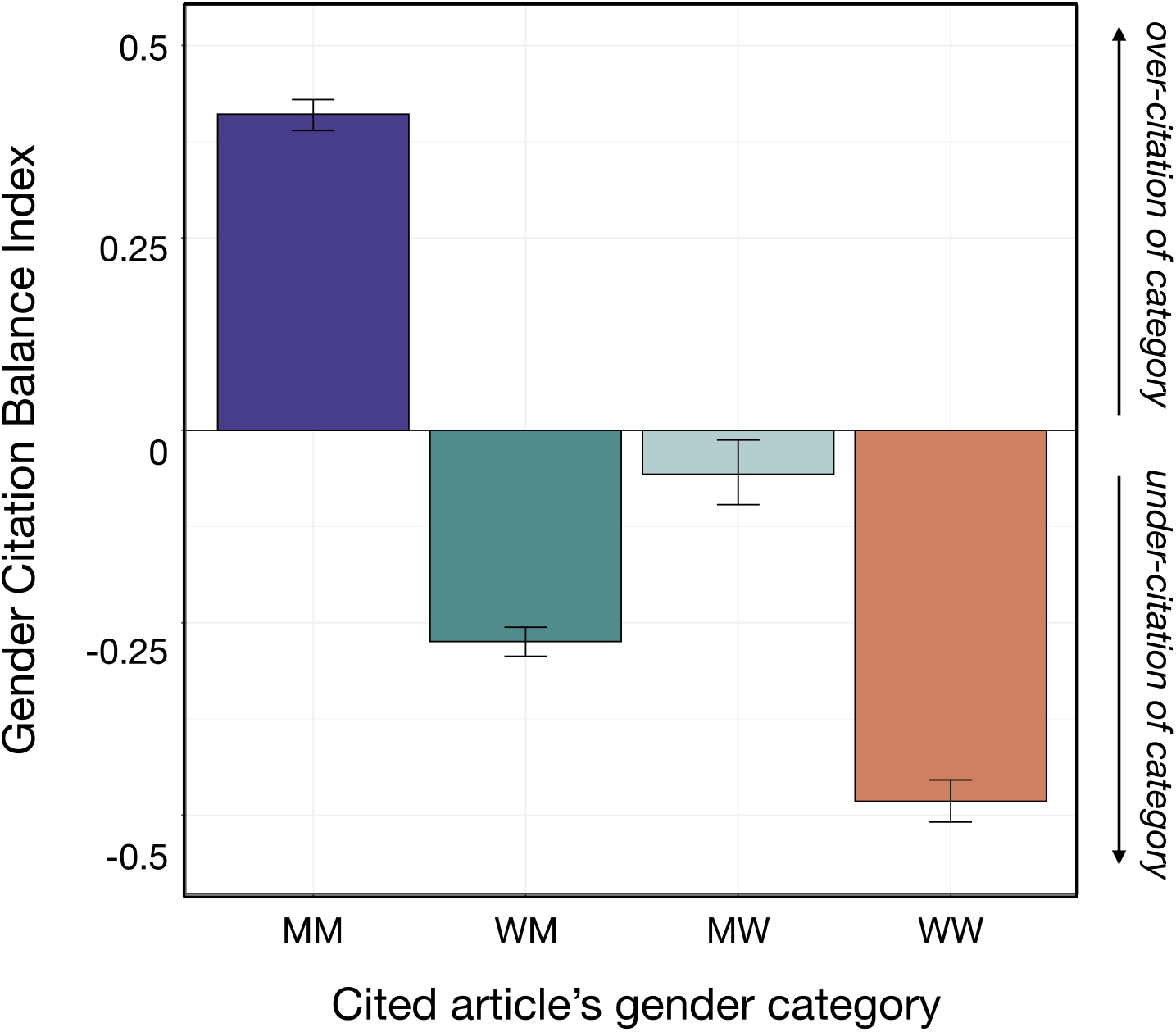
Gender Citation Balance Indices for the four gender categories of peer-reviewed articles published in *JoCN*. Error bars correspond to bootstrapped 95% confidence intervals.

**Figure 3.**
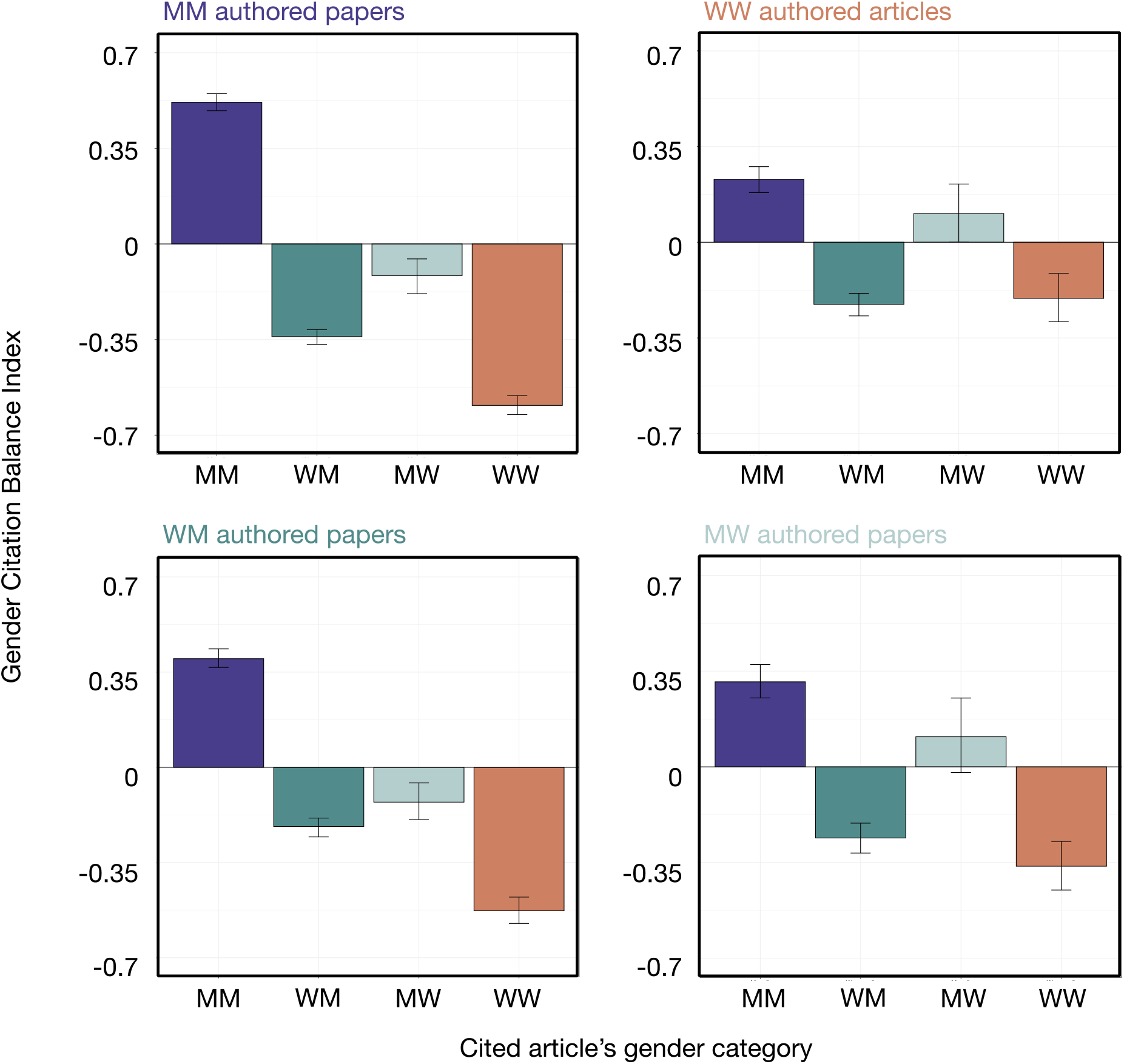
Gender Citation Balance Indices for peer-reviewed articles published in *JoCN*, broken down by citing articles’ gender category. Error bars correspond to bootstrapped 95% confidence intervals.

## DISCUSSION

We measured the degree to which the categorical gender balance of articles cited in the *Journal of Cognitive Neuroscience* from 2009-2020 reflected the gender balance of the authorship of the journal during that time frame. The results indicate that papers authored by men as the first and last authors have been over-cited compared to what would be expected based on the number of papers published by the journal that were authored by “MM” teams. By contrast, papers authored by teams with at least one woman in the first- and/or last-author position have been under-cited.

This finding indicates that the gender imbalance in citations previously reported for broad-scope neuroscience journals (Dworkin et al., 2020) extends to the sub-field of cognitive neuroscience. The fact that this pattern of imbalance is present in *JoCN* papers published by each of the four gender-defined groups that we have considered here (MM, WM, MW, WW) indicates that this imbalance results, at least in part, from systemic factors at play in the field overall.

In carrying out these analyses we deliberately limited ourselves to the “simple first step” (c.f., Dworkin et al., 2020) of quantifying and describing this phenomenon. Although we lack the expertise to propose specific interventions that may encourage prosocial behavior, it is our hope that this work contributes, in some modest way, to social norm messaging (Murrar et al., 2020) about the need to address inequities in the way that we carry out and communicate our science.

